# Integrating target capture with whole genome sequencing of recent and natural history collections to explain the phylogeography of wild-growing and cultivated *Cannabis*

**DOI:** 10.1101/2024.10.17.618884

**Authors:** Manica Balant, Daniel Vitales, Zhiqiang Wang, Zoltán Barina, Lin Fu, Tiangang Gao, Teresa Garnatje, Airy Gras, Muhammad Q. Hayat, Marine Oganesian, Jaume Pellicer, Alireza S. Salami, Alexey P. Seregin, Nina Stepanyan-Gandilyan, Nusrat Sultana, Shagdar Tsooj, Magsar Urgamal, Joan Vallès, Robin van Velzen, Lisa Pokorny

## Abstract

- *Cannabis* has provided important and versatile services to humans for millennia. Domestication and subsequent dispersal have resulted in various landraces and cultivars. Unravelling the phylogeography of this genus poses considerable challenges due to its complex history.
- We relied on a Hyb-Seq approach (combining target capture with shotgun sequencing), with the universal Angiosperms353 enrichment panel, to explore the genetic structure of wild-growing accessions and cultivars by implementing phylogenomic and population genomic workflows on the same Hyb-Seq data.
- Our findings support the treatment of *Cannabis* as a monotypic genus (*C. sativa* L.), structured into three main genetic groups—E Asia, Paleotropis, and Boreal—with clear phylogeographic signal despite significant levels of admixture. The E Asia group was sister to the Paleotropis and the Boreal groups. Individuals within the Paleotropis group could be further structured into three subgroups: Iranian Plateau, C & S China and Himalayas, and Indoafrica. Individuals from the Boreal group split into two subgroups: Eurosiberia and W Mongolia and Caucasus and Mediterranean. Hemp and drug-type landraces and cultivars consistently matched their putative geographic origin.
- These findings enhance our understanding of the genetic patterns in *Cannabis* and provide a framework for future research into its current and past genetic diversity.

## INTRODUCTION

*Cannabis sativa* L. (hereafter referred to as *Cannabis*) is one of the oldest multi-purpose crops, utilised by humans worldwide for thousands of years (Clarke & Merlin, 2013). It has been used as fibre (ropes, fabric, paper), medicinally (over 200 recorded uses), as food (nutrient-rich seeds), as well as in various magico-religious rituals (Balant *et al*., 2021a,b). Despite its long history of use, *Cannabis* was broadly deemed illegal at the beginning of the 20^th^ century, primarily because of its psychoactive properties. Consequently, studies on *Cannabis* became scarce and relied almost completely on hemp cultivars or on plant material confiscated by law enforcement. Nonetheless, spurred by recent legalization efforts, the *Cannabis* research and industry are now experiencing a revival in the agronomic, medicinal, and recreational sectors. Although there are several chromosome-level reference genomes and abundant whole genome sequencing (WGS) data available for *Cannabis*, these data predominantly originate from modern hemp cultivars or drug strains with unknown geographic origins and limited genetic diversity (e.g., van Bakel *et al*., 2011; Braich *et al*., 2020; Grassa *et al*., 2021; but see also Gao *et al*., 2020; Ren *et al*., 2021; Chen *et al*., 2022). Meanwhile, comprehensive studies including wild-growing and landrace individuals remain scant, which is why sampling these individuals across the entire natural distribution of this genus is much needed to better understand *Cannabis* genetic diversity and geographic structure (Kovalchuk *et al*., 2020).

*Cannabis* belongs to the Cannabaceae, an angiosperm family with ten genera and over 100 species (WFO, 2024). Within the family, two closely related species stand out for their economic significance: hops (*Humulus lupulus* L.), which plays a key role in the beer industry; and *Cannabis*, which is widely used in both medical and recreational sectors (Fu *et al*., 2023). *Cannabis* is a dioecious plant (except for some monoecious cultivars; Clarke & Merlin, 2013; Heer *et al*., 2024), typically a diploid (2n = 20; although natural triploids and tetraploids exist), with an average genome size of ∼1 pg/1C (Sharma *et al*., 2015; Balant *et al*., 2022; Philbrook *et al*., 2023).

Different centres of origin of the genus across Eurasia have been proposed, but palaeobotanical studies on subfossil pollen indicate that *Cannabis* most probably originated somewhere close to the NE Tibetan Plateau ∼27 million years ago (Mya) (Clarke & Merlin, 2013; McPartland *et al*., 2018, 2019; Zhang *et al*., 2018a; McPartland & Small, 2020). From there, it likely first spread west, reaching Europe approximately 6 Mya, and then east, arriving in E China around 1.2 Mya. Despite its current widespread use across India, the oldest subfossil pollen remains indicate that it reached the Indian subcontinent only ∼30 thousand years ago (Kya) (McPartland *et al*., 2019; Rull, 2022).

Similarly, the domestication of *Cannabis* has long been the subject of discussion. Some authors proposed a single C Asian domestication event (Schultes *et al*., 1974), whereas others suggested several independent ones (Vavilov, 1926; McPartland *et al*., 2018, 2019; Jin *et al*., 2021); however, the high concentration of early archaeological remains, together with the latest study by Ren *et al*. (2021), suggest that *Cannabis* was first domesticated in E Asia, approximately 12 Kya. Although it was initially cultivated as a multipurpose crop, selection for specific type-use cultivars might have started ∼4 Kya, leading to the development of separate ‘Hemp-type’ vs. ‘Drug-type’ plants (Ren *et al*., 2021). Since then, humans have been instrumental in *Cannabis* dispersal across C and E Asia, Europe, along the Himalayas, and on the Indian subcontinent. Subsequently, with the establishment of numerous trading routes, such as the Silk Road, and the expansion of multiple empires, human-mediated dispersal intensified across Eurasia and towards Africa, reaching the Americas with the European colonization and the Atlantic slave trade. Currently, dispersal in the opposite direction is happening and modern cultivars are being reintroduced into native areas, resulting in admixture with local landraces and wild-growing *Cannabis* populations (Abel, 1980; Clarke & Merlin, 2013).

The taxonomy of *Cannabis* has historically been complex, influenced by cultural biases and legal issues that led to confusion, with numerous synonyms inconsistently applied to taxa across different geographic regions (McPartland & Guy, 2017). The first known differentiation between European and Asian *Cannabis* was recorded by Ibn-al-Baitār ca. 1240 (Lozano Cámara, 2017; McPartland & Guy, 2017); however, it was not until the 18^th^ century that Linnaeus (*C. sativa*; 1753) and Lamarck (*C. indica* Lam.; 1783) scientifically described two distinct species. In the past two centuries, various taxonomic approaches based on genetics, morphology, and phytochemistry have been proposed, with several researchers treating *Cannabis* as a polytypic genus, identifying two or three species with various subspecies or varieties (Janischevsky, 1924; Vavilov, 1935-translated in 1992; Emboden, 1974; Schultes *et al*., 1974; Anderson, 1980; Clarke & Merlin, 2013; Jin *et al*., 2021). One of the first comprehensive studies, including a broad range of wild-grown and landrace *Cannabis* accessions with a worldwide distribution, was conducted by Hillig (2005a). Based on allozyme variation, morphological characters, and phytochemical profiles, he recognised two *Cannabis* species with six so-called ‘biotypes’: *C. sativa* for the accessions from the Levant, Europe, and N Asia (with hemp and feral ‘biotypes’) and *C. indica* for accessions from S, W, and E Asia, as well as Africa (with narrow-leaflet drug, wide-leaflet drug, hemp, and feral ‘biotypes’). He suggested a third species, *C. ruderalis* Janisch., might also exist; however, the sampling of individuals potentially belonging to this third putative species was too sparse to confirm its existence (Hillig, 2005b). Based on Hillig’s findings (2005a,b), Clarke and Merlin (2013) adopted a similar classification, with three species and six subspecies.

In contrast to this polytypic taxonomic concept, others considered *Cannabis* to be a monotypic genus, recognizing only *C. sativa* (Small & Cronquist, 1976; Sawler *et al*., 2015; Small, 2015; Lynch *et al*., 2016; McPartland *et al*., 2018; Ren *et al*., 2021; Lapierre *et al*., 2023), albeit with different infraspecific taxonomic divisions. McPartland & Small (2020), who follow the classification proposed by Small & Cronquist (1976) that recognises two subspecies within *C. sativa* (ssp. *sativa* and ssp. *indica*), carried out a large-scale revision of morphological traits, building on past genetic and phytochemical studies. Thus, within ssp. *indica*, they identified two domesticated (D) and two wild type (WT) varieties: var. *indica* (D) and var. *himalayensis* (WT) from S Asia, and var. *afghanica* (D) and var. *asperrima* (WT) from C Asia. The study by Ren *et al*. (2021), which mostly included hemp cultivars and drug strains, along with some wild-growing populations, also indicated that *Cannabis* should be considered as a single species, with individuals clustering into four genetic groups: ‘Basal cannabis’, ‘Hemp-type’, ‘Drug-type feral’, and ‘Drug-type’. Other WGS and microsatellite markers studies have also observed differentiation between geographic regions, and between hemp and drug accessions, sometimes with further distinctions within the drug genetic pool, identifying two separate groups (Sawler *et al*., 2015; Lynch *et al*., 2016; Schwabe & McGlaughlin, 2019; Woods *et al*., 2023). However, none of these studies included feral samples from either Mongolia or Africa, and they included few samples from the Caucasus, the Levant, and C & W Asia—areas otherwise reported as potentially very diverse (Soorni *et al*., 2017; McPartland & Small, 2020; Dehnavi *et al*., 2024). Moreover, several investigations analysing only within-country genetic diversity, found complex population structure within *Cannabis* in, e.g., China (Zhang *et al*., 2018a; Chen *et al*., 2022), USA (Busta *et al*., 2022), Iran (Soorni *et al*., 2017; Shams *et al*., 2020; Dehnavi *et al*., 2024), Morocco (Benkirane *et al*., 2024), and India (Pandey *et al*., 2023).

Based on cultivation purpose, morphology, and chemical composition, *Cannabis* plants can also be described as hemp-type (primarily grown for fibre and seed production) and drug-type, based on Δ9-tetrahydrocannabinol (THC) concentration (Hurgobin *et al*., 2021) or on the THC and cannabidiol (CBD) ratio (THC-dominant, balanced THC:CBD, and CBD dominant, that is, Type I, Type II, and Type III, respectively; Small & Beckstead, 1973). Outside of academic environments, drug-type plants are typically classified as ‘sativa’, ‘indica’, or ‘hybrid’ (McPartland & Guy, 2017); however, several studies have demonstrated that these informal classifications are not supported by genetic data (Sawler *et al*., 2015; Schwabe & McGlaughlin, 2019; Watts *et al*., 2021).

Recent studies have relied on high-throughput sequencing approaches such as genotyping-by-sequencing (GBS) or whole genome sequencing (WGS) to study *Cannabis*; however, no previous study has attempted to use a target-capture sequencing (TCS) approach to investigate the evolution of *Cannabis*. Furthermore, none of these studies has explored the potential of herbarium specimens, which could offer valuable insights into the past distribution of *Cannabis* genotypes. Hyb-Seq (Weitemier *et al*., 2014; Dodsworth *et al*., 2019), that is, TCS combined with low-coverage WGS, is an affordable method (Hale *et al*., 2020) proven very effective for sequencing not only recent and silica-dried tissue, but also historical collections (i.e., herbarium tissue), where DNA template is often highly degraded, which up until recently had thwarted their inclusion in genetic studies (Villaverde *et al*., 2018; Brewer *et al*., 2019; Shee *et al*., 2020). Different probe sets (TCS kits) for specific plant families (e.g., Asteraceae, Mandel *et al*., 2014; Euphorbiaceae, Villaverde *et al*., 2018; Dioscoreaceae, Soto Gomez *et al*., 2019) or larger taxonomic groups (e.g., flagellate land plants, Breinholt *et al*., 2021) have been developed. The universal Angiosperms353 enrichment panel is a probe set which includes 353 orthologous nuclear protein-coding genes found in single copy across all flowering plants (Johnson *et al*., 2019). Although originally conceived to study phylogenetic relationships above the species level, it has successfully been used for population-level analyses of various flowering plant groups (Slimp *et al*., 2021; Wenzell *et al*., 2021; Beck *et al*., 2021; Yardeni *et al*., 2022; Crowl *et al*., 2022; Phang *et al*., 2023), as well as domesticated landraces (Van Andel *et al*., 2019).

To address the taxonomic inconsistencies and to gain a clearer understanding of the genetic structure of *Cannabis*, we conducted a comprehensive sampling (with emphasis on wild-growing populations) focusing on its native distribution range (taking special care to include individuals from previously under-sampled areas). Relying on the same Hyb-Seq dataset, we carried out phylogenomic analyses to clarify the taxonomic status of wild-growing and landrace *Cannabis* accessions, and we implemented population genomics analyses to better understand how populations are structured. In this manner, we linked macro- and microevolutionary scales to shed light on the phylogeography of *Cannabis*.

## MATERIAL AND METHODS

### Sampling and Molecular Protocols

For the ingroup, we sampled 94 *Cannabis sativa* L. individuals with emphasis on populations across Eurasia (Fig. 1). Fifty-eight samples were obtained from living plants, dried in silica gel, and 36 samples were secured from herbaria. For the outgroup, three *Humulus scandens* and three *H. lupulus* SRAs, corresponding to WGS and RNA-sequencing data, were downloaded from the NCBI repository (see Table S1 for details).

**Fig. 1.**
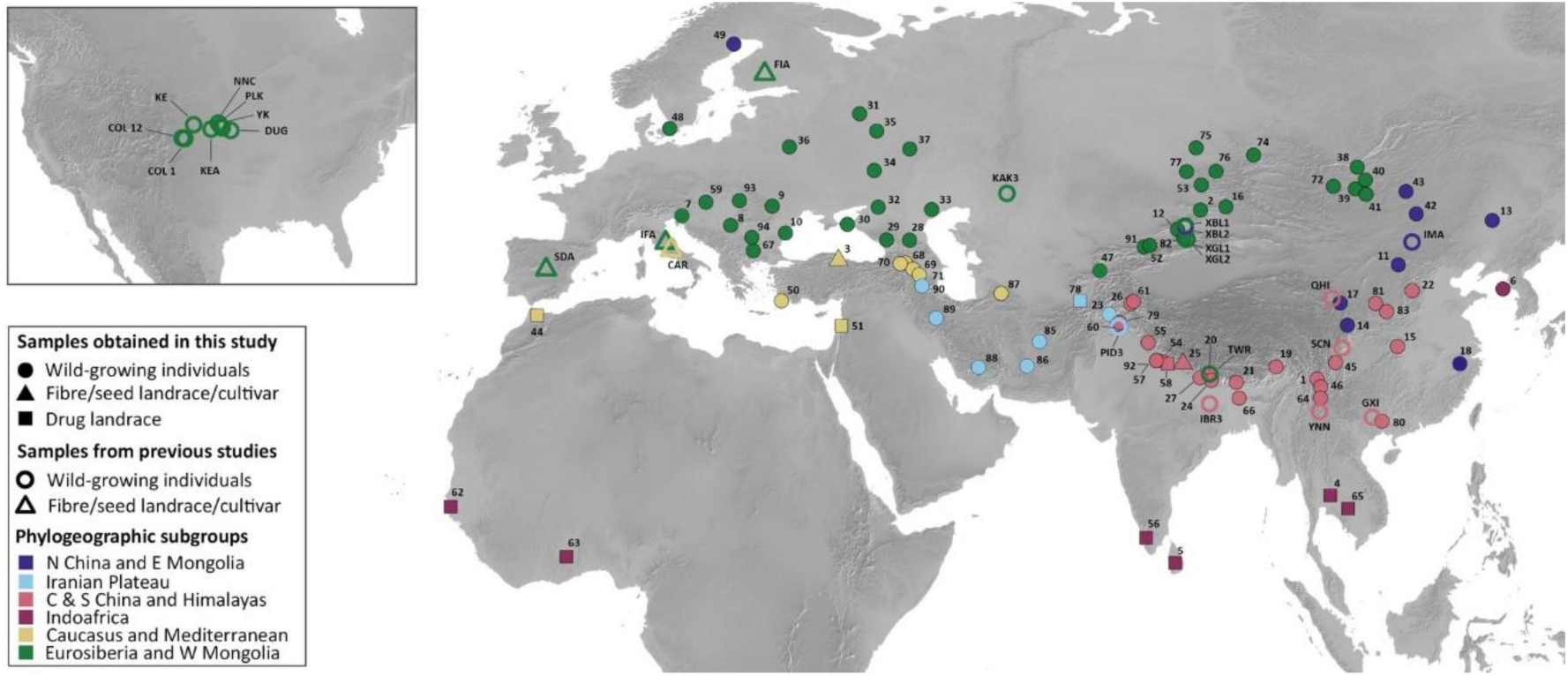
Geographic distribution of samples included in this study, with individuals coloured according to the subgroups obtained in the phylogenomic analysis (see Fig. 2). The shapes indicate *Cannabis* accession types, them being, wild-growing (circles), fibre/seed (triangles), and drug (squares) types. Additionally, filled shapes are newly analysed Hyb-Seq samples, while empty shapes are NCBI SRAs corresponding to WGS data mined for our Hyb-Seq targets. The inset shows USA wild-growing populations mined from NCBI SRAs. Drug cultivars mined are not shown. For more detailed information see Supplementary Table S1. The map was made with *Natural Earth* (Free vector and raster map data @ naturalearthdata.com).

DNA of 94 *Cannabis* individuals was extracted either using the E.Z.N.A. SP Plant DNA Kit (Omega Bio-Tek, Norcross, GA, USA) or a modified CTAB protocol (Doyle & Doyle, 1987). DNA concentration was measured with a Qubit fluorometer (Thermo Fisher Scientific, Waltham, MA, USA) using dsDNA BR Qubit assays. The extractions yielded on average 2,000 ng of DNA.

DNA extractions were sent to Daicel Arbor Biosciences (Ann Arbor, MI, USA), who provide target capture sequencing services (myReads®). They carried out DNA quantitation, genomic library preparation (with dual indexing), target enrichment (nine libraries per capture reaction), and Illumina® sequencing. Captures were performed following the myBaits v5.03 protocol, using the myBaits® Expert Angiosperms353 enrichment panel (Johnson *et al*., 2019), with an overnight hybridization and washes at 65° C. Enriched libraries were then pooled in approximately equimolar ratios, alongside the original genomic libraries at a ratio of 75% enriched to 25% original genomic libraries. Samples were sequenced on an Illumina® NovaSeq 6000 platform on a partial S4 PE150 lane, resulting in an approximate 108 Gbp total.

### Sequencing Data Processing

The de-multiplexed raw sequences were first filtered and trimmed using fastp v0.23.4 (Chen, 2023), removing adapters and low-quality reads (-f 20 -t 5 -F 20 -T 5 -g -x -W 3 -r -M 20 -q 20 -l 40 --detect_adapter_for_pe; for lower quality samples flags -q 15 and -l 30 were used instead), and checked with FastQC (Andrews, 2010) and MultiQC (Ewels *et al*., 2016) before and after filtering with fastp.

HybPiper v2.1.6 (Johnson *et al*., 2016) was then used to recover the single-copy nuclear genes from the Angiosperms353 enrichment panel with the target file mega353.fasta (McLay *et al*., 2021) and the *assemble* flag and the ‘bwa’ option. We checked for potential paralogues using the *paralog_retriever* flag, calculated statistics with the *stats* flag, and visualised the gene recovery using the *recovery_heatmap* flag. Due either to the presence of paralogues (73_RUS_SB) or because of extremely low coverage (84_CHN_ANH), we eliminated two individuals from further analysis. The *max_overlap* script (Shee *et al*., 2020) was then used to calculate a coverage score for each of the remaining accessions and sequences. Four more *Cannabis* individuals (12_CHN_XIN, 15_CHN_HUB, 17_CHN_QIN, and 18_CHN_ZHN), three *Humulus* accessions (DRR024392, SRR24774240, and SRR24774242), and eight genes (6514, 6886, 6705, 6893, 6713, 6565, 6557, and 5354) were also eliminated to reduce noise and remove underrepresented, incomplete, and unevenly distributed sequences across accessions from our data matrix. The supercontigs (exons plus flanking regions) of 345 target genes for the 91 remaining individuals (88 *Cannabis* and three *Humulus* individuals) were then retrieved using the *retrieve_sequences* flag selecting the ‘supercontig’ option.

### Nuclear Species Tree Inference

Retrieved sequences were then aligned with MAFFT v7.520 (Katoh & Standley, 2013) (using flag *auto*). Exploratory gene trees were constructed with FastTree 2 v2.1.11 (Price *et al*., 2010) and TreeShrink v1.2.1 (Mai & Mirarab, 2018) was used to automatically prune outlier branches, using the false positive tolerance rate (α) of 0.05 and the ‘per-species’ option. The output was then re-aligned using MAFFT (same settings as above) and trimmed with trimAl v1.4.1 (Capella-Gutiérrez *et al*., 2009) using relaxed settings (gap threshold set to 0.3, while keeping at least 30% of the original alignment).

The gene trees were inferred under maximum likelihood (ML) with IQ-TREE v2.2.6 (Nguyen *et al*., 2015) using ModelFinder to select the best fit DNA substitution model (Kalyaanamoorthy *et al*., 2017) and choosing the non-parametric Shimodaira–Hasegawa approximate likelihood ratio tests (SH-aLRT; Guindon *et al*., 2010) for assessing branch support values with 1,000 replicates. For the resulting ML gene trees, unsupported branches were collapsed using the ‘nw_ed’ tool from the Newick Utilities v1.6.0 package (Junier & Zdobnov, 2010) with threshold 0% SH-aLRT, as recommended by Simmons & Gatesy (2021). The coalescent species tree was then inferred using ASTRAL-III (Zhang *et al*., 2018b) and, since branch lengths in the resulting topology come in coalescent units, RAxML-NG v 1.2.1 (Kozlov *et al*., 2019) was used (with flag *evaluate*) to estimate branch lengths in substitutions per site (pre-requisite for some of our downstream analyses). Gene tree vs. species tree incongruence was visualised with the *AstralPlane* package (Hutter, 2021) in R v4.3.2 (R Core Team, 2022), using the *astralProjection* function to plot quartet scores calculated in ASTRAL-III (using the ‘-t 2’ option) as pie charts. Trees were visualised in FigTree v1.4.4 (Rambaut, 2018).

### Phylogenomic Placement of WGS Accessions

We downloaded 64 publicly available *Cannabis sativa* WGS SRAs from the NCBI repository (see Suppl. Table S1 for details), which we then placed in our nuclear species tree. Using fastp (Chen, 2023), these raw sequences were also quality-filtered and trimmed (with flags -f 15 -t 5 -F 15 -T 5 -g -x -W 3 -r -M 20 -q 20 -l 40 --detect_adapter_for_pe; additionally, and to prevent batch effects, alternative quality filters were also used, i.e., -f 20 -t 7 -F 20 -T 7), and quality-checked with FastQC (Andrews, 2010) and MultiQC (Ewels *et al*., 2016). The four previously eliminated samples (12_CHN_XIN, 15_CHN_HUB, 17_CHN_QIN, and 18_CHN_ZHN) were added to the 64 NCBI SRAs.

To recover the Angiosperms353 target genes from these WGS SRAs, we also used HybPiper (Johnson *et al*., 2016), following the same steps described above. No paralogues were found in the downloaded dataset; however, due to the diverse approaches (e.g., varying levels of sequencing depth) implemented by the different research teams who produced and shared their *Cannabis* WGS data, many samples had poor target gene recovery (to be expected, given that our 353 targets mostly appear in single-copy in the nucleus). We discarded 32 individuals that had < 200 target genes with sequences with < 50% of the mean target length, as well as the same eight genes flagged by the abovementioned *max_overlap* script (see Table S2 for details). As a result, we were left with 36 accessions, for which we retrieved supercontigs (exons and flanking regions) of 345 target genes using the *retrieve_sequences* flag and the ‘supercontig’ option in HybPiper.

These supercontigs were then aligned using the 91-individual alignment above as a constraint in MAFFT (Katoh & Standley, 2013) (with flags *add* and *keeplength*). The resulting alignments were pruned to extract the 36 accessions, which we then placed in the 91-individual ASTRAL species tree (following branch-length recalculation with flag *evaluate* in RAxML-NG, see above) with EPA-ng v0.3.8 (Barbera *et al*., 2019), using the best.Model file previously obtained with RAxML-NG (also with flag *evaluate*). The placement output was converted with GAPPA (Czech *et al*., 2020), using the function *guppy tog*, and visualised in FigTree.

### SNP Calling and Population Genomics Analyses

For SNP calling, we used the supercontig sequences (exons and flanking regions) of the 88 *Cannabis* individuals that were also included in the phylogenomic analyses. To call the SNPs, we followed the workflow designed by Slimp *et al*. (2021), with minor modifications. In brief, we generated a combined reference sequence from the longest supercontig recovered for each of the Angiosperms353 target genes. The variant detection was carried out with GATK4 v4.5.0.0 (McKenna *et al*., 2010). We refined the combined SNP data matrix using filters ‘QD < 5.0’, ‘FS > 60.0’, ‘MQ < 40.0’, ‘MQRankSum < −12.5’, and ‘ReadPosRankSum < −8.0’, with flag *missing-values-evaluate-as-failing*. Only SNPs that passed all filters above were then processed using BCFtools v1.20 (Danecek *et al*., 2021) and VCFtools v0.1.16 (Danecek *et al*., 2011) to eliminate multi-allelic variants, and to only keep SNPs with minimum 30% quality, minimum and maximum mean depth of 10 and 200, respectively, maximum missingness of 10%, and minor allele frequency of at least 10%. All individuals had coverage < 36 and > 40 missingness. Using PLINK v1.9 (Purcell *et al*., 2007), we additionally filtered the SNPs based on linkage disequilibrium, with settings --indep 50 5 2. On this fully filtered and unlinked SNP data matrix we calculated eigenvalues and eigenvectors for 20 principal component analysis (PCA) axes, also with PLINK.

The analysis of population structure was first carried out with STRUCTURE v.2.3.4 (Pritchard *et al*., 2000), as implemented in the ipyrad toolkit v0.9.52 (Eaton & Overcast, 2020). The filtered VCF file was first converted into a HDF5 file, with a linkage block size of one. We assigned individuals into six population groups, which matched the subgroups in our ASTRAL species tree, and then ran the analysis with burnin length one million and three million replicates. A range of K values (2–10) was tested in five independent runs and the most likely number of clusters was selected by detecting the highest values of the ΔK statistic (Evanno *et al*., 2005). The population structure was visualised as an ancestry matrix using the *geom_bar* function from the *ggplot2* package in R, and pie charts were projected onto a map obtained from *Natural Earth* with MAPMIXTURE package in R (Jenkins, 2024). Heterozygosity and pairwise identity-by-descent were calculated using PLINK. The fixation index (F_ST_) was calculated for each pair of the previously detected six phylogenetic groups using PLINK v.2.0 (Chang *et al*., 2015), and following the Hudson method (Bhatia *et al*., 2013).

## RESULTS

### Phylogenomic analyses reveal geographically defined groups

We used the Hyb-Seq approach to sequence 94 *Cannabis* individuals from across its entire native distribution (Fig. 1). Target enrichment with the Angiosperms353 universal probe set was successful for both silica-dried and herbarium samples in all but one individual. On average we obtained more than 18 million reads per individual, with ∼20% reads on target for silica-dried tissue and ∼23% for herbarium samples. Using the HybPiper ‘supercontig’ option, gene recovery rate was very high (median value for genes with at least 50% targeted gene-length recovered was 336 for silica-dried tissue and 334 for herbarium samples). We detected only six genes with putative paralogues, present in a single individual that was eliminated from downstream analyses. Finally, 88 *Cannabis* individuals (ingroup) and three *Humulus* accessions (outgroup) were included in the final dataset used to infer a species tree under the multispecies coalescent (MSC) theoretical framework (Figs. 2A & S1). Because WGS is not targeted, HybPiper retrieval was less efficient for downloaded WGS data for which, despite the high number of reads per sample (average > 96 million reads), on average only 0.42% reads mapped to the Angiosperms353 targets (with values ranging between 0.1% and 2.4%). Detailed information on target recovery statistics and max_overlap outputs can be found in Supplementary Tables S2 to S7.

**Fig. 2.**
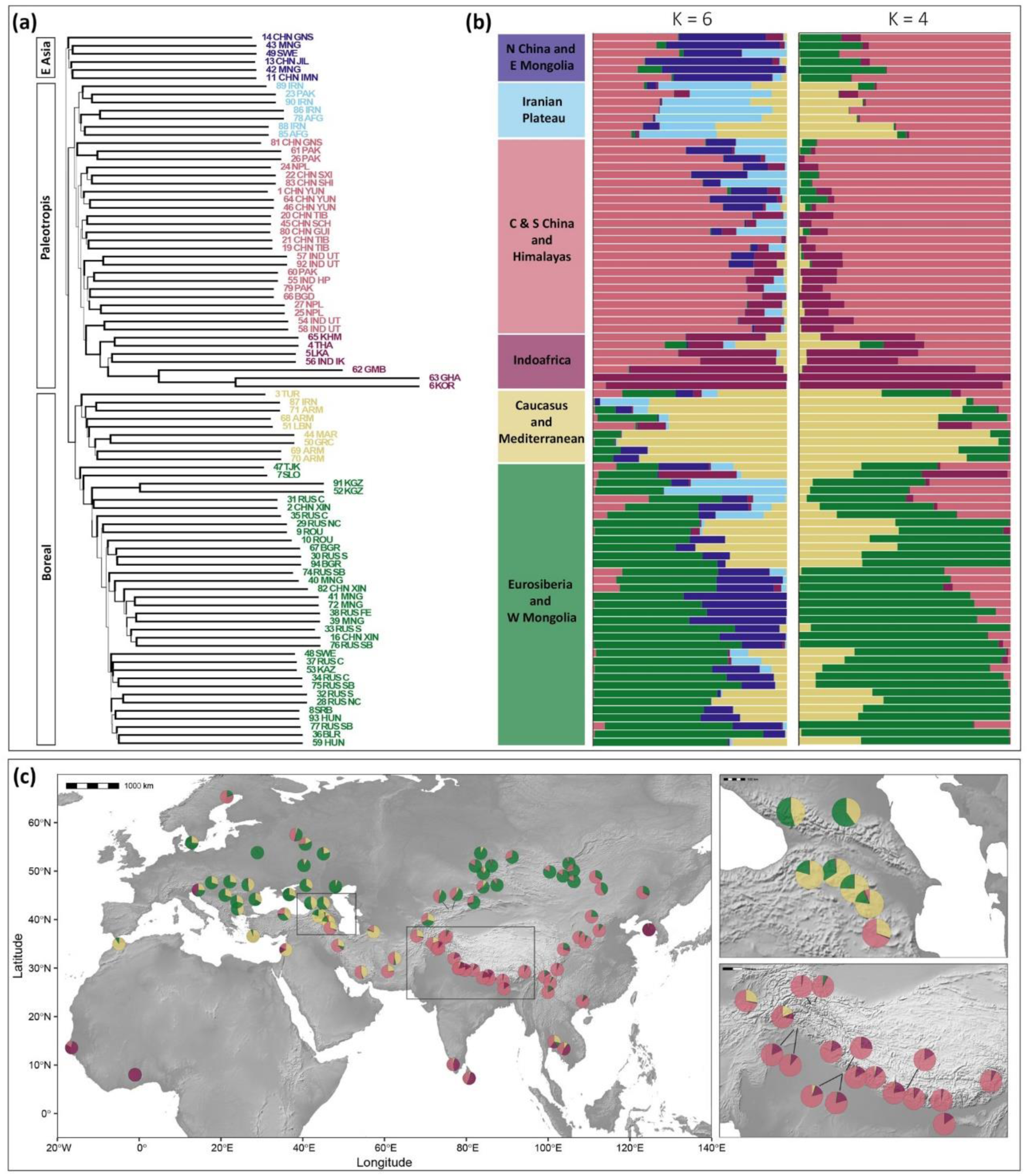
Phylogenomic and population genomic analyses reveal the complex genetic structure of *Cannabis sativa* (a) ASTRAL-III nuclear species tree (for topology with the outgroup see Supp. Fig. S1) inferred from 345 ML gene trees (estimated with IQ-TREE2 from filtered MAFFT alignments), showing three main groups (E Asia, Paleotropis, and Boreal) subdivided into six subgroups matching the geographic distribution of the samples analysed (only the 88 highest-quality samples shown). Branch thickness in the species tree is proportional to support measured as local posterior probabilities (LPP), and branch length is shown in coalescent units. (b) Admixture plots estimated in STRUCTURE from 2,875 (filtered and unlinked) SNPs called from the same 345 nuclear ortholog targets used to estimate the species tree. We show genetic admixture plots for the two most likely clustering scenarios (K = 4 and K = 6, as per ΔK statistic values). (c) Geographic distribution of samples coloured for K=4, with two insets zooming into the Caucasus and the Himalayas. The map was made with *Natural Earth* (Free vector and raster map data @ naturalearthdata.com).

The MSC species tree inferred from 345 nuclear gene trees clustered all *Cannabis* individuals together in a clade sister to genus *Humulus* (LPP = 1.0; Fig. S1). Within *Cannabis*, a division into three main genetic groups was observed. While local posterior support for these three main groups was very low, the individuals comprising them clustered into geographically distinctive subgroups (Figs. 2B & S1). The first group (E Asia group), which is sister to the other two main groups, consists of individuals from N China (provinces of Jilin, Gansu, and Inner Mongolia) and E Mongolia. All other *Cannabis* individuals belonged to either of the remaining monophyletic groups. The second group (Boreal group) consists of individuals predominantly present at latitudes above 40° N and the third group (Paleotropis group) of individuals from lower latitudes. The Paleotropis group can be further divided into the Iranian Plateau subgroup (a poorly supported group sister to all other Paleotropis individuals that includes accessions from Pakistan, Afghanistan, and Iran), and the C & S China and Himalayas subgroup (from China, Nepal, Bangladesh, N India and N Pakistan). This latter subgroup further extends into S India, Sri Lanka, W Africa, and SE Asia, forming a distinctive, highly supported, Indoafrica subgroup that is well-nested within the C & S China and Himalayas subgroup. On the other hand, the Boreal group is divided into two subgroups, them being the Caucasus and Mediterranean subgroup (from Armenia, Turkey, Greece, Lebanon, and Morocco) and the Eurosiberia and W Mongolia subgroup (from Europe, Russia, Kazakhstan, NW China, Kyrgyzstan, Tajikistan, and W Mongolia), which are reciprocally monophyletic, albeit with low support.

No batch effects were observed with regards to the placement of the 32 downloaded WGS SRAs and 4 newly sequenced individuals (with lower target capture success) into the existing MSC species tree. Instead, their placement matched the geographic origin of the samples, and not their use (i.e., hemp-type vs. drug-type; Figs. 1, 3 & S1, S3). The phylogenetically placed samples display longer branches (measured in substitutions per site) than those already present in the MSC species tree, which we attribute to an artefact resulting from their higher proportion of missing data.

Including the newly sequenced individuals and the downloaded WGS SRAs, most of these samples were collected from wild-growing plants (101), but we also incorporated seven hemp cultivars, one high CBD cultivar, and 16 other drug strains. As previously stated, rather than by their use type, samples matched their geographic origin. Thus, the Carmagnola (CAR) hemp cultivar and an unnamed hemp cultivar from Turkey (3_TUR) were nested within wild-growing plants of the Caucasus and Mediterranean subgroup, while the Finola (FIA), Fibranova (IFA), Delta Llosa (SDA), and Fedora (FED) hemp cultivars fell in the Eurosiberia and W Mongolia subgroup, both within the Boreal group. A multipurpose landrace from Nepal (25_NPL) primarily used for fibre production was nested in the C & S China & Himalayas subgroup, within the Paleotropis group. As for the drug types, many were placed in the Paleotropis group; Haze drug strain (HAE) and landraces from Thailand (4_THA), Cambodia (65_KHM), Sri Lanka (7_LKA), S India (56_IND_IK), and W Africa (62_GMB & 63_GHA) all belonged to the Indoafrica subgroup, while N India (54_IND_UT & 58_IND_UT) drug landraces were nested in the C & S China and Himalayas subgroup. Drug strains Ruderalis indica (RIA), Hindu Kush (HKH), Purple Kush (PPK), Top 44 (TOP), and Afghanistan landrace (78_AFG) were all nested in the Iranian Plateau subgroup. However, drug landraces from Morocco (44_MAR) and Lebanon (51_LBN) belonged to the Caucasus and Mediterranean subgroup, nested within the Boreal group. Lastly, the high CBD cultivar (CBDRx) was placed within the Iranian Plateau subgroup, in the Paleotropis group.

### Population genomic analyses reveal extensive admixture across the native range

Using the longest supercontig sequences (exons and flanking regions) per target gene for the variant mapping, we were able to recover a total of 68,212 single nucleotide polymorphisms (SNPs), from the 88 *Cannabis* accessions also included in our phylogenomic workflow. Of these SNPs, 2,875 passed our robust filtering settings and were used for downstream population structure analyses.

The PCA of the filtered and unlinked SNP dataset confirmed the geographical signal revealed by the phylogenomic analyses (Fig. 4). The two first PCA axes confirm the separation of the three main groups recovered in the MSC nuclear tree. There is also clear clustering of individuals following the subgroups these main groups are divided into in the nuclear species tree (Fig. 1), which is consistent throughout different PCA axes (Fig. 4). Together, the first two axes explain ∼30% of the total variation (PC1: 20.46%; PC2: 9.56%). *Cannabis* individuals are thus structured into a Eurosiberia and E Mongolia subgroup (top-centre left), a Caucasus and Mediterranean subgroup (bottom centre), a N China and E Mongolia subgroup (top centre), an Iranian Plateau subgroup (middle), a C & S China and Himalayas subgroup (top right), and a Indoafrica subgroup (bottom right); with some exceptions (i.e., 3_TUR, 4_THA, 7_LKA, 51_LBN; Fig. 4A). The third axis (PC3) explains 8.32% of the variance and places the Indoafrica subgroup and the Caucasus and Mediterranean subgroup at opposite ends of a continuum, with most other individuals clustering in the middle (Fig. 4B & 4C).

The STRUCTURE analyses (Fig. 2B) indicated that the most optimal number of clusters is four (Fig. S3). This clustering scheme is apparently inconsistent with the three main groups divided into six subgroups we observe in our nuclear species tree (Fig. 2A). Interestingly, some of these six subgroups are characterised by specific admixture patterns, also evident when we map the geographic distribution of the individuals’ ancestry (Fig. 2C). Indeed, when increasing the cluster number to K=6 (second most likely clustering scheme), the specific admixture pattern characterising these six subgroups comes to the foreground (Fig. 2B). For instance, the N China and E Mongolia subgroup is a mixture of two clusters (K=4, salmon and green; K=6, salmon and dark blue). The same could be said of the Iranian Plateau subgroup (K=4, salmon and mustard; K=6, salmon and mustard with light blue); however, for this latter subgroup, the predominant cluster when K=6 (light blue) is barely found elsewhere (except for the Kyrgyzstan individuals, 52_KGZ & 91_KGZ). Meanwhile, the Indoafrica subgroup is mostly composed of a single cluster (maroon), with barely any hints in most of its individuals of the predominant cluster in the Paleotropis group (salmon), where this subgroup is otherwise nested. Similarly, the Caucasus and Mediterranean subgroup is mostly composed of a single cluster (mustard, Fig. 4), with barely a touch of the predominant cluster in the Boreal group (green), where this subgroup belongs. However, this latter predominant cluster does characterize the Eurosiberia and W Mongolia subgroup (green), regardless of the clustering scheme.

There are some individuals (i.e., 3_TUR, 4_THA, 7_ SLO and 47_TJK) with noticeably admixed ancestry profiles for either clustering scheme (measured as inbreeding coefficient, F). The genetic structure of the Turkish hemp cultivar for example is showing a highly admixed profile with high outbreeding (F_03_TUR_ = −0.522), and points to recent admixture with genetically distant individuals from different genetic backgrounds. The Slovenian sample, found growing wild near a field, likely resulted from a recent unintentional crossing between a nearby drug strain and a hemp cultivar, as evidenced by its high outbreeding (F_7_SLO_ = −0.483). The herbarium sample 47_TJK (collected from a wheat field in Tajikistan, back in 1969) shows mixed genetic ancestry, but it does not present too high outbreeding (F_47_TJK_ = 0.071). On the opposite end, we can find some individuals (i.e., 6_KOR, 19_CHN_TIB, 21_CHN_TIB, 26_PAK, 33_RUS_S, 36_BLR, 44_MAR, 50_GRC, 61_PAK, and 63_GHA) barely showing any admixture at all. While the mean inbreeding coefficient for the highly admixed samples was generally low (F = −0.358), the latter unmixed samples show relatively high mean inbreeding coefficient (F = 0.276), compared to the mean F value of 0.083 for all the samples (Table S8). As expected, the Lebanese (F_51_LBN_ = 0.2677) and Moroccan (F_44_MAR_ = 0.2717) drug landraces exhibit high inbreeding.

With regards to the inbreeding coefficient of the six subgroups identified in the nuclear species tree, the Indoafrica subgroup had the highest mean value (F = 0.133), while the Iranian Plateau had the lowest mean value (F = −0.005; Table S8). To check if the phylogenetic subgroups existing in close geographic proximity also shared the most genetic diversity, we calculated the fixation index (F_ST_) between them. The lowest genetic distance was found between the N China and E Mongolia subgroup and the C & S China and Himalayas subgroup (F_ST_ = 0.036), and the highest between the Eurosiberia and W Mongolia subgroup and the Indoafrica subgroup (F_ST_ = 0.155; Table 1). Samples from the Indoafrica subgroup in general exhibit the highest genetic divergence from other phylogeographic subgroups and show high levels of inbreeding (F > 0.2). Only two samples in this subgroup show high outbreeding (5_LKA and 4_THA; F < −0.2), which may be due to recent hybridization for landrace improvement, as suggested by their genetic admixture profiles. Notably, the sample with the highest outbreeding (F_4_THA_ = −0.498) appeared roughly in the middle of our PCA (PC1 through PC3 axes), most distant to all other Indoafrican samples (label 4, Fig. 4). The highest proportion of shared alleles was found between individuals 6_KOR and 63_GHA (0.791), while the average value was 0.023 (Table S9).

**Table 1.** Pairwise fixation index (Hudson F_ST_) values between phylogeographic subgroups.

| Phylogeographic subgroup pairs | | Hudson $F_{ST}$ |
| --- | --- | --- |
| Eurosiberia and W Mongolia | Indoafrica | 0.155 |
| Caucasus and Mediterranean | Indoafrica | 0.136 |
| N China and E Mongolia | Indoafrica | 0.126 |
| Indoafrica | Iranian Plateau | 0.120 |
| Indoafrica | C & S China and Himalayas | 0.090 |
| Caucasus and Mediterranean | N China and E Mongolia | 0.086 |
| Eurosiberia and W Mongolia | C & S China and Himalayas | 0.080 |
| Eurosiberia and W Mongolia | Iranian Plateau | 0.077 |
| Caucasus and Mediterranean | C & S China and Himalayas | 0.076 |
| Caucasus and Mediterranean | Eurosiberia and W Mongolia | 0.061 |
| N China and E Mongolia | Iranian Plateau | 0.060 |
| N China and E Mongolia | Eurosiberia and W Mongolia | 0.058 |
| Caucasus and Mediterranean | Iranian Plateau | 0.048 |
| Iranian Plateau | C & S China and Himalayas | 0.039 |
| N China and E Mongolia | C & S China and Himalayas | 0.036 |

## DISCUSSION

The classification of genus *Cannabis* has historically been subject to numerous interpretations, ranging from multiple species to just one. To shed light on the taxonomic status of *Cannabis*, we analysed a comprehensive set of 88 *Cannabis* wild-growing and landrace individuals across its natural distribution area (Figs. 1, S1 & S2), filling in previous sampling gaps (Levant, Caucasus, C & W Asia, and Mongolia). We relied both on recently collected silica-dried tissue samples and on historical herbarium materials. Consistent with previous work (Ren *et al*., 2021), our MSC species tree (inferred from 345 single-copy nuclear orthologs; Fig. 2A), as well as the results from our population genomics analyses, fully support *Cannabis* as a monotypic genus (*C. sativa*, LPP = 1.0; Fig. S1) sister to the hops genus (*Humulus*). We detected admixture and low genetic differentiation even among the most distantly related populations.

### *Cannabis sativa* Phylogeographic Structure

Within *C. sativa*, we observe three geographically well-defined groups (Figs. 2A & S1), where the E Asia group is sister to the Paleotropis and the Boreal groups. These geographic groups broadly agree with the findings of previous genetic studies (Hillig, 2005b; Ren *et al*., 2021; Fig. S4). However, contrary to what Ren *et al*. (2021) found, our groups match the geographic distribution of individuals and not the use type, even when we place their WGS data into our nuclear species tree (Figs. 3 & S3). Additionally, we further subdivide the Paleotropis and Boreal groups into three and two subgroups, respectively, albeit with low support. A weakly supported backbone topology has also been inferred for other Angiosperms353 population-level studies (e.g., *Castilleja*, Orobanchaceae; Wenzell *et al*., 2021), where considerable conflict among gene trees was observed. Indeed, we also detect extensive gene-tree conflict in *Cannabis* (see quartet scores for a selection of branches in Fig. S1). This conflict could stem from introgression or deep coalescence (shared ancestral alleles), as hinted in our population admixture plots (Fig. 2B). To generate these admixture plots in STRUCTURE, we used 2,875 filtered and unlinked SNPs obtained from 345 nuclear orthologs (comprising exons and their flanking regions), from both fresh silica-dried material and herbarium tissue samples. Although exons from the Angiosperms353 targets are relatively conserved across land plants, the non-coding flanking regions provided sufficient variability to uncover consistent phylogeographic patterns within *Cannabis* populations. Granted that WGS data (Ren *et al*., 2021; Woods *et al*., 2023) does result in abundant SNPs that may capture greater variability, it does not necessarily resolve problematic nodes, as support values in Ren *et al*. (2021) indicate.

**Fig. 3.**
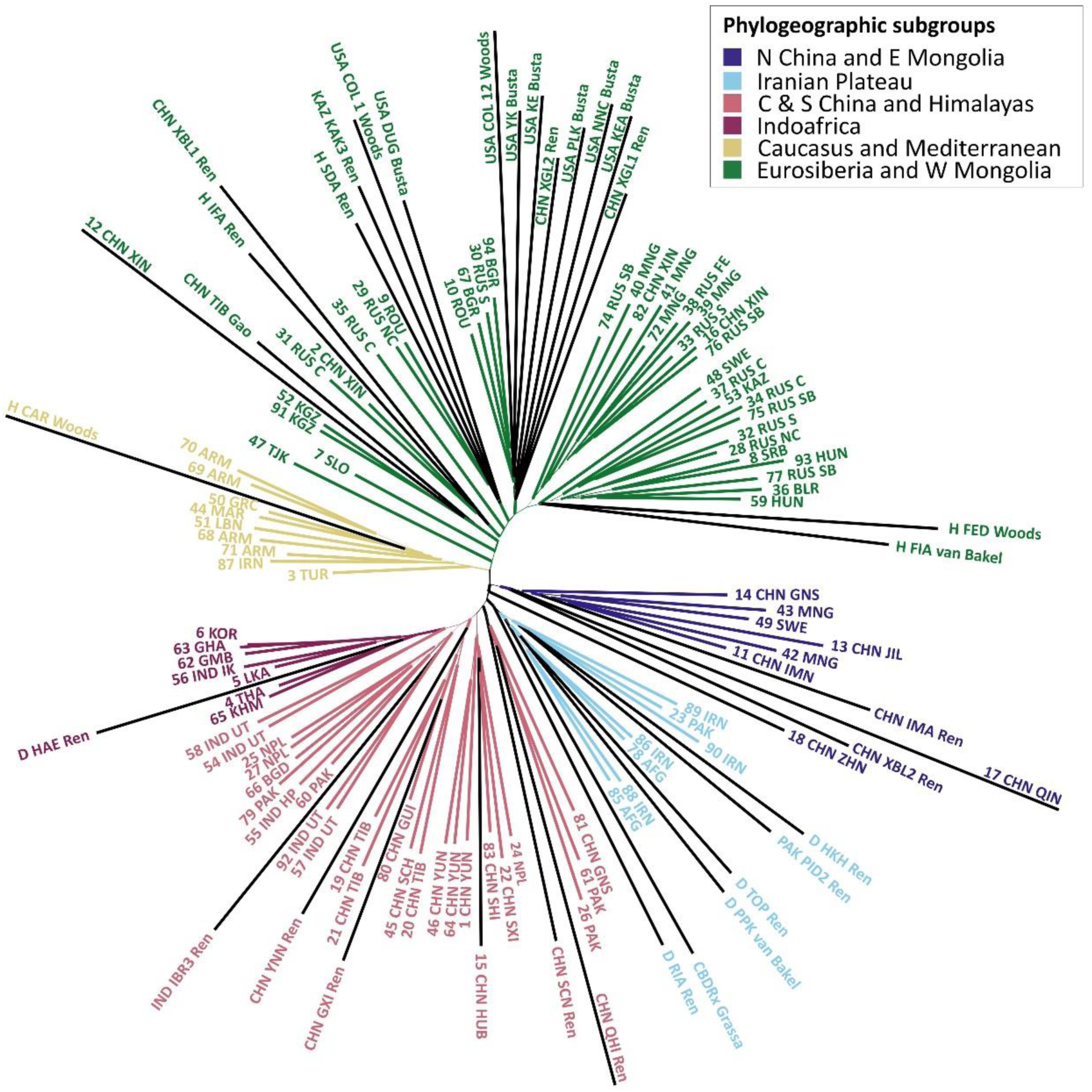
Unrooted topology depicting the phylogenomic placement (done with EPA-ng) of the four lower-quality Hyb-Seq samples and the 32 WGS samples downloaded from the NCBI SRA database (black terminal branches) into the nuclear species tree inferred from 345 nuclear targets for *Cannabis* (for rooted topology with the outgroup see Supp. Fig. S2).

**Fig. 4.**
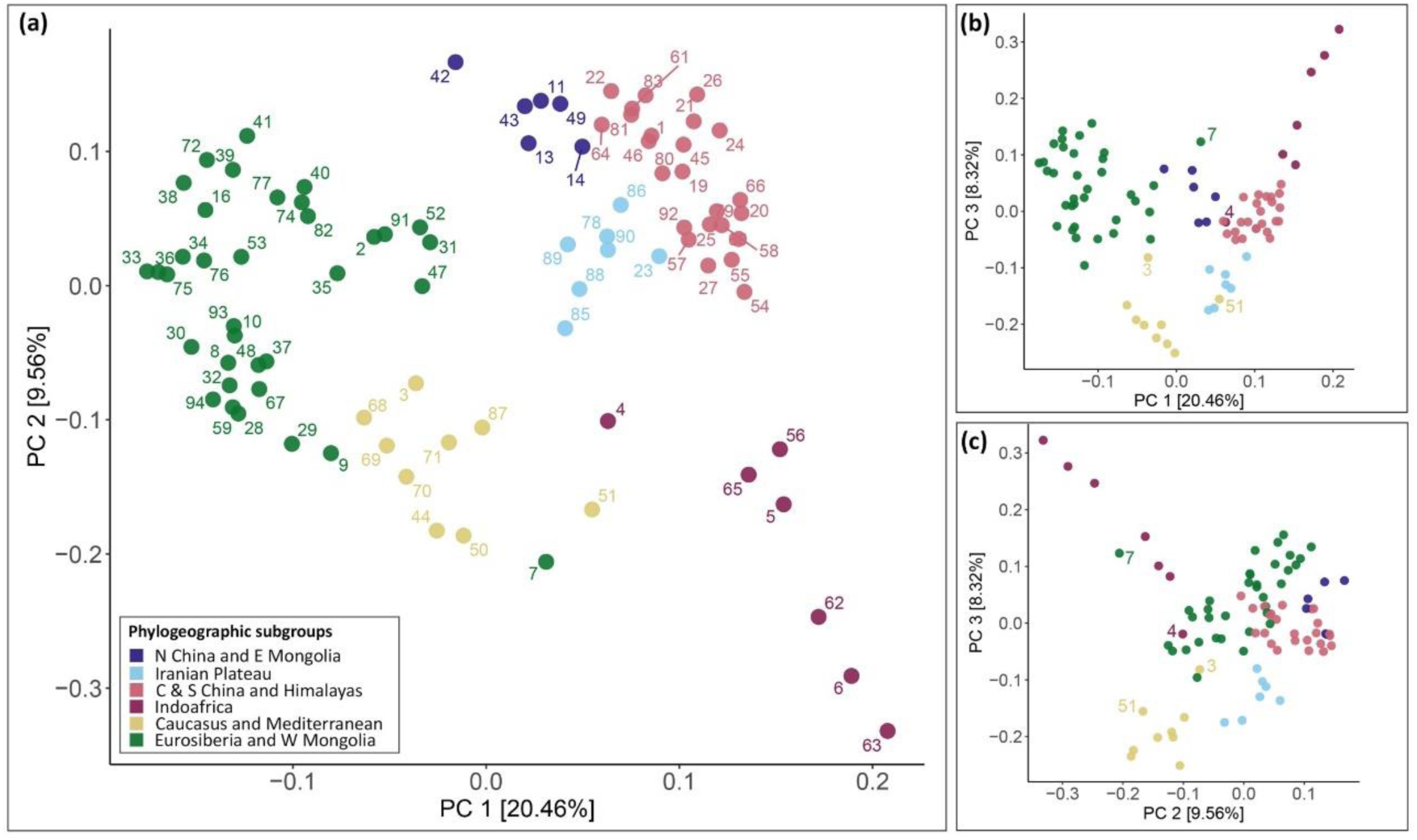
Principal component analysis (PCA; done with PLINK) of *Cannabis sativa* individuals for the 2,875 (filtered and unlinked) single nucleotide polymorphisms (SNPs) called from the same 345 nuclear ortholog targets (comprising exons and their flanking regions) used to estimate the nuclear species tree (supercontig data matrix). Colours correspond to the six phylogeographic subgroups identified in the phylogenomic analysis (see Fig. 2). (a) First and second PCA axes. (b) First and third PCA axes. (c) Second and third PCA axes.

### The E Asia Group

The E Asia group comprises samples from N China and E Mongolia (hence the subgroup name). Interestingly, an individual from N Sweden also clustered within this subgroup. Given it was collected nearby a church, it is plausible this plant represents a lineage introduced by Swedish missionaries working in China in the 19^th^ and 20^th^ centuries (Gregersen, 2023). One could argue that the most recent common ancestor of *C. sativa* might have originated close to this N China and E Mongolia region (which would be in agreement with previous palaeobotanical findings; McPartland *et al*., 2019) and, from there, spread westwards into the Altai Mountains (giving rise to the Boreal group), and south-westward into the Hengduan Shan and the Himalayas (resulting in the Paleotropis group), to later intersect in the axis formed by the C Asian Tien-Shan, Pamir-Alay, and Hindu-Kush mountain ranges.

Out of the six ‘Basal cannabis’ accessions from Ren *et al*. (2021) that we could include in our study, only two fell in our E Asia group (Figs. 3, S3 & S5). This discrepancy may result from varying levels of missing data in their dataset versus ours or it could be attributed to molecular methodological differences, i.e., WGS is anonymous (orthology needs to be assessed) and Hyb-Seq is targeted (orthology is known), where anonymous sequencing could introduce excessive noise, while targeted sequencing could lack sufficient signal (Fuentes-Pardo & Ruzzante, 2017; Dodsworth *et al*., 2019). Additionally, as suggested by Halpin-McCormick *et al*. (2024), sampling biases could be driving the results instead. Our study focused on wild-growing individuals and landraces, with special attention to previously underrepresented areas across the entire distribution (e.g., Levant, Caucasus, and Mongolia), whereas the study by Ren *et al*. (2021) predominantly sampled commercial hemp cultivars and drug strains.

### The Paleotropis Group

Within the Paleotropis group, we identified three distinct subgroups. The first one, the Iranian Plateau subgroup, includes wild-growing and drug-type plants from Iran, Afghanistan, and Pakistan. These samples exhibit a specific admixture pattern characterising its two adjacent subgroups (Fig. 2C). Prior to our study, Iranian samples had not been extensively included in *Cannabis* phylogeographic analyses. Soorni *et al*. (2017) identified two distinct groups within Iranian *Cannabis* populations: one comprised of plants from W Iran and another from E Iran. This differentiation may be influenced by admixture between W Iranian accessions and the Caucasus and Mediterranean subgroup, as evidenced by the distinct patterns observed in our results (Fig. 2).

The second subgroup in the Paleotropis group, the C & S China and Himalayas subgroup, includes *Cannabis* plants with a distribution ranging from Pakistan to C China, spreading across the Himalayas, the Hengduan, and even the Qinling mountains. It encompasses both hemp- and drug- type plants, as well as wild-growing plants.

Nested within C & S China and Himalayas subgroup is a third one we denominate the Indoafrica subgroup, which is primarily composed of drug-type plants typically found at latitude ca. 10° N. A comparable subgroup was also found in a study by Lynch *et al*. (2016). While Hillig (2005b) did not find distinctions between individuals from Iran, Afghanistan, and Pakistan compared to other Asian samples, he did note that samples from sub-Saharan Africa and SE Asia clustered together within what he called the ‘*indica* gene pool’, which has a distribution like that of our Paleotropis group (Fig. S4). Interestingly, a sample from an island off the coast of S Korea (6_KOR) also belongs to our Indoafrica subgroup; however, as it was collected from a ruderal environment, it is possible that it represents an escaped individual with a genotype closely related to those found in W Africa.

Numerous other studies have consistently identified two subgroups within drug-type *Cannabis* accessions, commonly referred to as ‘sativa’ and ‘indica’ (Sawler *et al*., 2015; Lynch *et al*., 2016; Schwabe & McGlaughlin, 2019; Vergara *et al*., 2021). In our phylogenomic placement analyses (Figs. 3 & S3), we included five drug strains (van Bakel *et al*., 2011; Ren *et al*., 2021) and one high CBD cultivar (Grassa *et al*., 2021). The strain named Haze grouped with samples from the Indoafrica subgroup, likely reflecting its development from landraces primarily originating from Thailand and S India (Clarke & Merlin, 2013). The remaining samples (CBDRx, HKH, PPK, TOP, and RIA) were placed within the Iranian Plateau subgroup and are generally believed to stem from landraces from Pakistan and Afghanistan, which subsequently have been extensively crossbred (Clarke & Merlin, 2013). Put simply, drug-type accessions (be them cultivated or wild-growing) are most closely related to the populations where they might have originated. It is therefore likely that these subgroups (Indoafrican versus Iranian Plateau) represent separate ancestral gene pools that eventually gave rise to the so-called ‘indica’ and ‘sativa’ drug types. Nowadays, these distinctions have been blurred due to crossing and inconsistent labelling practices, resulting in low reliability in the naming of *Cannabis* drug strains (Sawler *et al*., 2015; Schwabe & McGlaughlin, 2019; Watts *et al*., 2021).

### The Boreal Group

The Boreal group spans from NW Africa and Europe, across Russia into C Asia and even W Mongolia (Figs. 1 & 2). Within this group, we identified two subgroups: the Caucasus and Mediterranean subgroup and the Eurosiberia and W Mongolia subgroup. Previous studies did not detect the former subgroup, likely due to limited sampling in the region. The few studied samples until now were usually classified under what Hillig (2005b) denominated the ‘*sativa* gene pool’ (polytypic framework).

The Caucasus and Mediterranean subgroup encompasses hemp cultivars from Turkey and Italy, drug landraces from Morocco and Lebanon, and wild-growing samples from Iran, Armenia, and Greece. The distinctive admixture pattern the Lebanese drug landrace exhibits seems to corroborate Clarke’s (1998) assertion regarding the introduction of germplasm from India. Clarke & Merlin (2013) proposed a S Asian origin for Moroccan landraces, but our study supports a Caucasus/Levantine origin. Onofri *et al*. (2015) identified shared SNP mutations between Moroccan and certain Afghan landraces, maybe indicating a putative shared genetic ancestry with the Iranian Plateau subgroup through the Caucasus and Mediterranean subgroup (see mustard for K = 6 in Fig. 2B). ‘Sieved hashish’ production (resin that is collected from dried *Cannabis* plants and filtered through several sieves, to separate and collect trichomes rich in cannabinoids and terpenes) has been documented in Pakistan, Afghanistan, and Iran (Iranian Plateau subgroup). Additionally, Morocco, Lebanon, Turkey, and Greece, all areas where the genotype of the Caucasus and Mediterranean subgroup predominates, are also known for their ‘sieved hashish’ production (Abel, 1980; Clarke, 1998). It is therefore plausible that both *Cannabis* and the knowledge of ‘sieved hashish’ production spread across the Mediterranean region through traders from the Caucasus/Levant, who themselves could have acquired that knowledge from C Asian peoples.

The second subgroup in the Boreal group, Eurosiberia and W Mongolia, covers an extensive area and aligns well with Hillig’s (2005b) ‘*sativa* gene pool’ (polytypic framework). It primarily consists of wild-growing plants from Europe, Russia, C Asia, NE China, and W Mongolia, but it also includes all North American individuals phylogenomically placed in our species tree (Figs. 1 inset, 4 & S3), partially in agreement with previous studies (Ren *et al*., 2021; Busta *et al*., 2022). The presence of C Asian plants in this subgroup is surprising, since they are morphologically different from typical *Cannabis* plants growing in Europe (Vavilov, 1926; Clarke & Merlin, 2013). Nonetheless, Ren *et al*. (2021) found that a sample from Uzbekistan aligned with hemp-type plants, which is consistent with our findings. Samples from NW China, Tajikistan, and Kyrgyzstan exhibit unique admixture patterns, often blending the Iranian Plateau subgroup with the Eurosiberia and W Mongolia subgroup. The samples from Kyrgyzstan show specific genetic structure not observed in other samples (Fig. 2B), with a possible introgression from Iranian Plateau populations. Historically, this region was known for high-quality ‘sieved hashish’ production by Muslim Uyghurs, although its production in the region was banned by the Soviets and Chinese in the 19^th^ and 20^th^ centuries, who instead introduced hemp cultivars promoting fibre production (McPartland & Small, 2020 Supp. Mat. 1).

This C Asian region is a melting pot of mountain ranges (Tien-Shan, Pamir-Alay, Hindu-Kush), biogeographic regions (Palearctic and Indomalaya), and cultures that seems to harbour genetically and morphologically diverse *Cannabis* plants in close proximity, which is surprising for a wind pollinated plant with no obvious reproductive barriers (McPartland & Small, 2020; Halpin-McCormick *et al*., 2024). Our results are in agreement with McPartland & Small (2020), that made a comprehensive revision of over one thousand herbarium specimens and identified the Pamir-Alay and Hindu-Kush mountains as a contact zone for different *Cannabis* genetic pools. Isolation of the Iranian Plateau subgroup, C & S China and Himalayas subgroup, and the Eurosiberia and W Mongolia subgroup was possibly maintained by inaccessible mountainous terrain and/or different cultural practices. In the northern areas of Eurasia drug use was secondary and plants were selected primarily for fibre use and seed production (with some exceptions, e.g., extensive hashish production in the Turpan region in NW China). In southern Eurasian regions *Cannabis* was more frequently consumed for its psychoactive properties (Clarke & Merlin, 2013). While ‘sieved hashish’ is favoured in Arab countries (Iranian Plateau and Caucasus and Mediterranean subgroups), ‘charas’ (hand-rubbed resin from living *Cannabis* plants) and ‘ganja’ (smoking of dried female inflorescences) are preferred in Hindu countries (C & S China and Himalayas subgroup). Clarke & Merlin (2013) proposed that plants in dry climates were selected for ‘sieved hashish’ production, favouring trichomes that fall off easily. Conversely, in the humid and rainy Himalayas loose trichomes would be disadvantageous, leading to the selection of traits that allow trichomes to withstand the rain and the higher humidity levels. ‘Sieved hashish’ production in these areas was therefore replaced by production of hand-rubbed *charas* or by directly smoking the dried female inflorescences (‘ganja’). These different traits reflect distinct genetic subgroups that are well-suited to their respective environments; the human-driven selection for different drug production styles may have further aided genetic isolation despite their geographic proximity.

### Concluding Remarks

Using both phylogenomic and population genomic approaches, we have gained deeper insights into the genetic patterns in *Cannabis*. Our results support the taxonomic treatment of *Cannabis* as a single species, *Cannabis sativa*, with three main groups (E Asia, Paleotropis, and Boreal) and six subgroups (see above). Unlike some previous studies, our findings show that individuals group according to their geographical distribution rather than their use type (e.g., hemp vs. drug-type).

Hyb-Seq has shown success in dealing with older and degraded herbarium specimens (Dodsworth *et al*., 2019), with some studies including herbarium specimens dating back to the 19^th^ century (Villaverde *et al*., 2018; Brewer *et al*., 2019; Shee *et al*., 2020; Gardner *et al*., 2022; Moreyra *et al*., 2023). Our study demonstrated that *Cannabis* herbarium specimens can be successfully analysed and integrated in population genomics studies. This opens the possibility of using historical herbarium specimens, collected before modern germplasm exchange and widespread crossing obscured the population structure of wild *Cannabis* populations, to illuminate the past distribution of this species and potentially detect some unquestionably wild specimens.

Since the Angiosperms353 probe set targets relatively conserved genes across land plants, it may lack the resolution needed for more detailed population genomics analyses in *Cannabis*. While WGS offers higher resolution and would allow for better integration with existing datasets, it does not perform well with older herbarium specimens. The lack of resolution could be addressed by designing a Cannabaceae specific probe set which could integrate the Angiosperms353 targets with other low-to-single-copy genes (e.g., those shared across Rosids that have successfully been used to infer phylogenomic relationships in Cannabaceae; Fu *et al*., 2023) and further refined by incorporating, functional genes of agronomic interest, such as those involved in cannabinoid biosynthesis or in fibre development (as previously done for the yam family (Dioscoreaceae); Soto Gomez *et al*., 2019). This combined strategy would offer a more comprehensive tool for future *Cannabis* research. Additionally, morphometric analysis (e.g., using leaves, as proposed by Balant *et al*., 2024) and phytochemical characterization could further clarify group delimitation within *Cannabis*.

## Acknowledgements

We would like to thank Joan Uriach Marsal (Uriach Laboratories), CIJA Preservations SL, and Cannaflos – Gesellschaft für medizinisches Cannabis mbH for additional financial support, and CIJA Preservations SL for providing valuable seed accessions. We would also like to thank Madison Bullock, Ciprian C. Buzna, Marco Calvi, David Criado, Teodora Dalmacija, Branko Dolinar, İrem Erdoğan, Pol Fernandez, Nikolai Friesen, Sònia Garcia, Dmitry German, Peter Glasnović, María Luisa Gutiérrez Merino, Oriane Hidalgo, Neus Ibáñez, Matthew G. Johnson, Bing Liu, Jordi Lopez, Kunigunda Macalik, Marina Olonova, Alexandra Papamichail, Sonja Siljak-Yakovlev, Vladimir Stevanović, Boštjan Surina, Paul Szatmari, Vladimir Vladimirov, Yushi Ye, and several herbaria (ALTB, CL, IBSC, KP, LD, MW, PE, and SNUA) and their curators. This research was supported by projects WECANN (CGL2017-84297-R, Ministerio de Ciencia, Innovacion y Universidades), Generalitat de Catalunya (CLT051, 2017SGR1116, and 2021SGR00315), Institut d’Estudis Catalans (PRO2020-2024-S02-VALLES). MB benefited from FPI predoctoral contract funded by the Ministerio de Ciencia, Innovacion y Universidades (PRE2018-083226). LP benefited from a Ramón y Cajal grant (RYC2021-034942-I) funded by MCIN/AEI/10.13039/501100011033 and by the European Union “NextGenerationEU”/PRTR. AG benefited from a postdoctoral grant of the Universitat de Barcelona funded by NextGeneration EU funds (Margarita Salas 2022–2024). We acknowledge support of the publication fee by the CSIC Open Access Publication Support Initiative through its Unit of Information Resources for Research (URICI). This research project was made possible through the access granted by the Galician Supercomputing Center (CESGA) to its supercomputing infrastructure. The supercomputer FinisTerrae III and its permanent data storage system have been funded by the Spanish Ministry of Science and Innovation, the Galician Government, and the European Regional Development Fund (ERDF).

## Author contributions

MB, TGar, DV, and LP conceptualized the study, with contributions from RvV regarding the population genomics analyses. TGar, JV, and JP secured the funding. MB, ZB, BD, LF, TGao, AG, MQH, MO, ASS, APS, NS-G, NS, ST, MU, JV, and ZW conducted fieldwork or provided key herbarium specimens. MB did molecular lab work. MB, DV, LP, and RvV designed the analytical workflow, which MB and LP used to process the data. DV, TGar, LP, ZW, and RvV supervised the work. MB and LP wrote the first draft of the paper, which all authors reviewed, commented on, and edited.

## Data availability

The raw Hyb-Seq data are available on NCBI Sequence Read Archive (SRA) under BioProject PRJNA1162815. The additional publicly available sequences used in the study are listed in Supplementary Table S1. The intermediate files (trimmed alignments, gene and species trees, filtered VCF file and structure and admixture outputs) and workflow are available from Zenodo (to add).

