## Supplementary material for "Integrating target capture with whole genome sequencing of recent and natural history collections to explain the phylogeography of wild-growing and cultivated *Cannabis*": Fig. S2

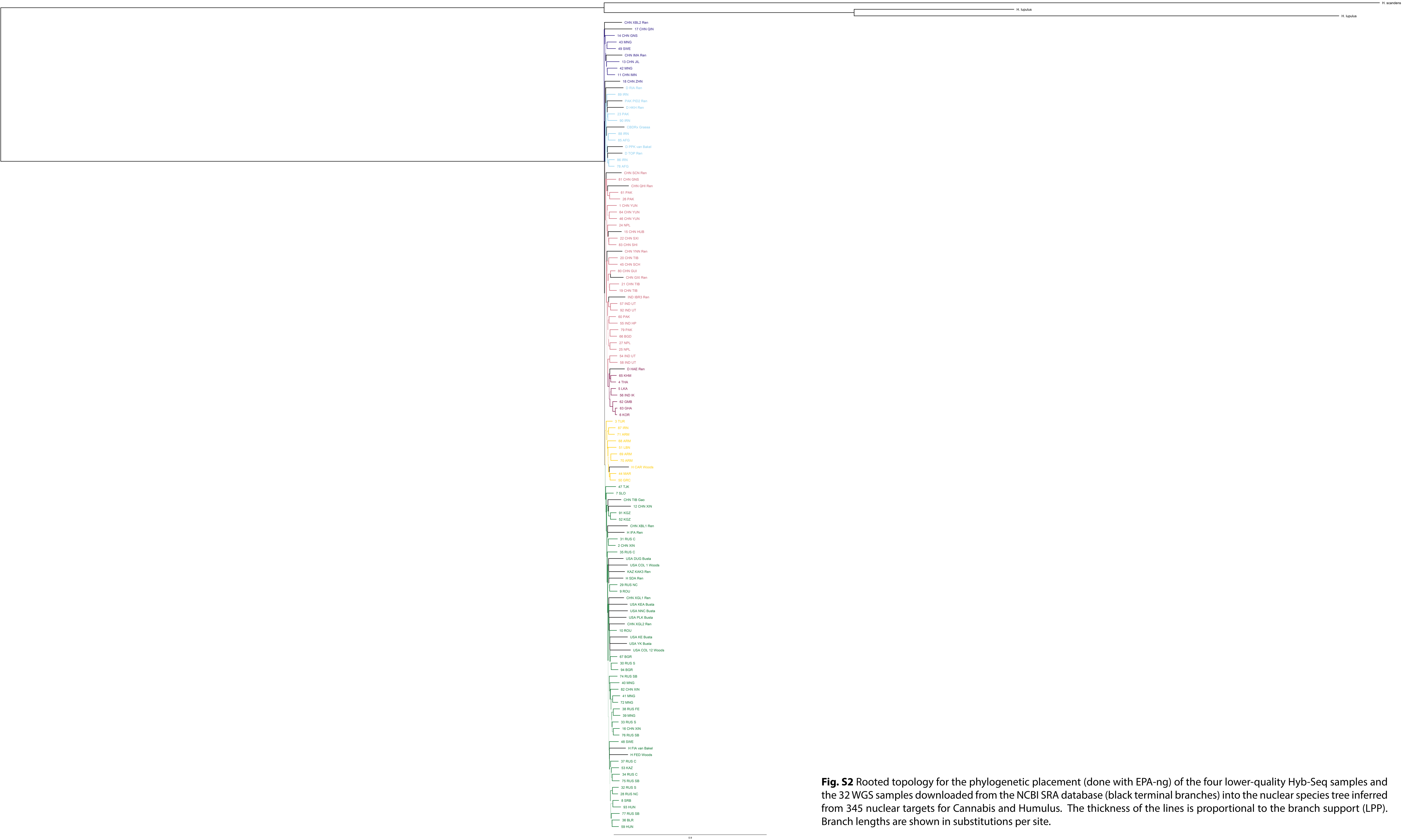


**Fig. S2** Rooted topology for the phylogenetic placement (done with EPA-ng) of the four lower-quality Hyb-Seq samples and the 32 WGS samples downloaded from the NCBI SRA database (black terminal branches) into the nuclear species tree inferred from 345 nuclear targets for Cannabis and Humulus. The thickness of the lines is proportional to the branch support (LPP). Branch lengths are shown in substitutions per site.
