## Supplementary material for "Integrating target capture with whole genome sequencing of recent and natural history collections to explain the phylogeography of wild-growing and cultivated *Cannabis*": Figs S1, S3, S4

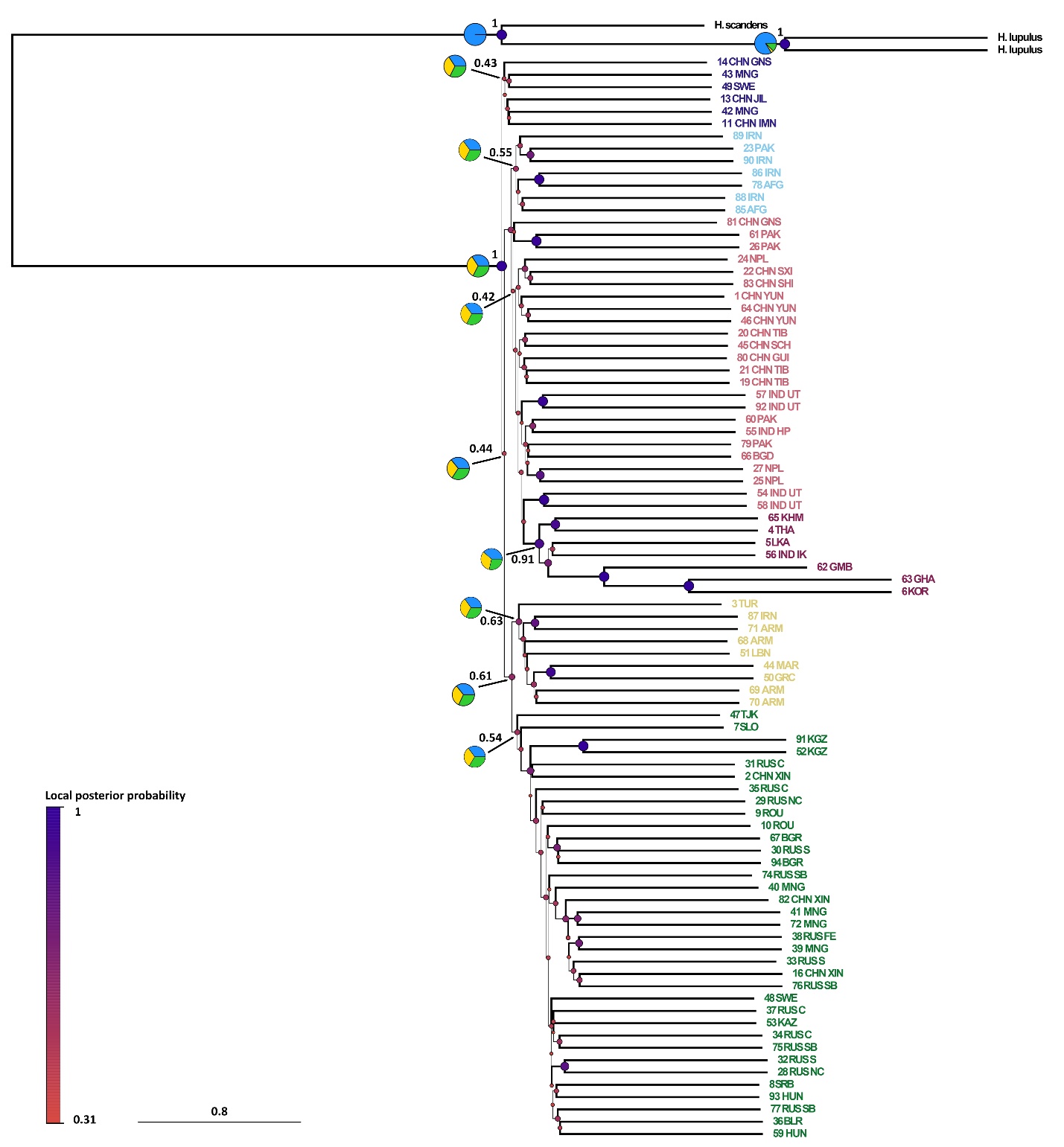

**Fig. S1** ASTRAL-III nuclear species tree inferred from 345 ML genes trees (estimated with IQ-TREE2 from filtered MAFFT alignments), showing the outgroup (Humulus sp.) and three main Cannabis groups (E Asia, Paleotropis, and Boreal) subdivided into six subgroups matching the geographic distribution of the analysed samples. Pie charts show gene tree conflict at the nodes of the six subgroups as calculated in ASTRAL-III and visualised with the AstralPlane package in R. For these same branches, the values next to pie charts indicate local posterior probabilities (LPP), while for the remaining branches, the thickness of the lines is proportional to the branch support (LPP). Branch lengths are shown in coalescent units.

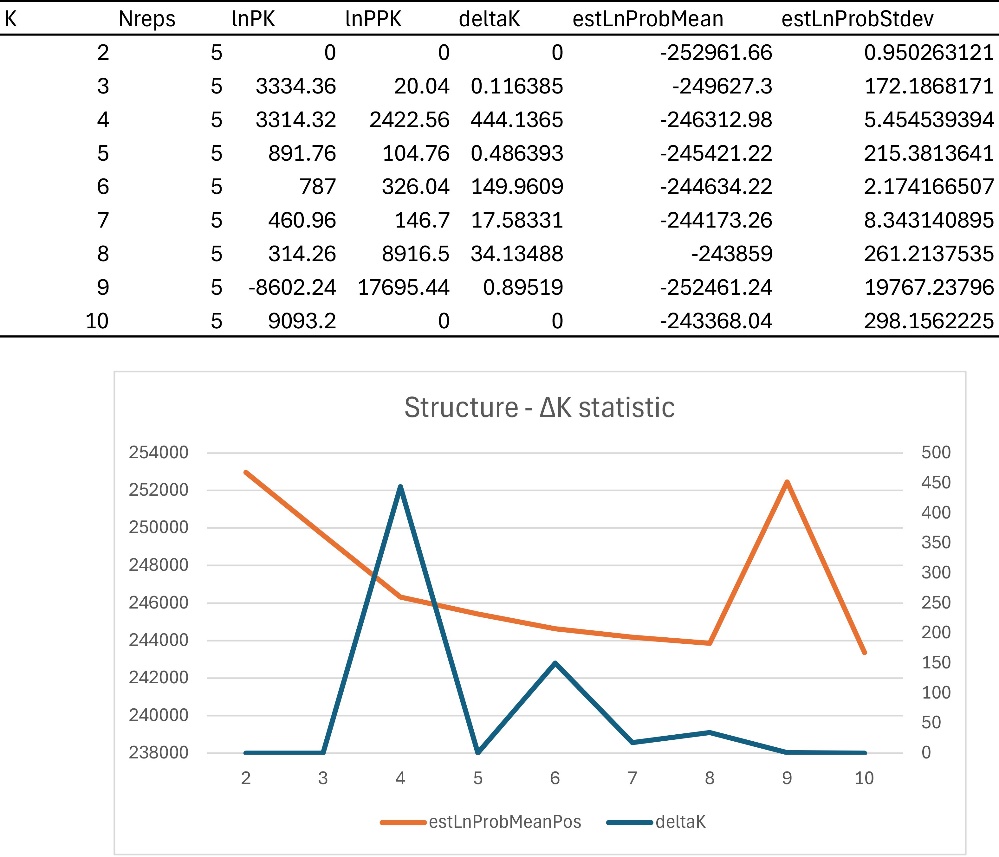

**Fig. S3** ΔK plot (blue) and mean for estimated log probability of data (orange) of all five STRUCTURE independent runs for K = 2 to K = 10.

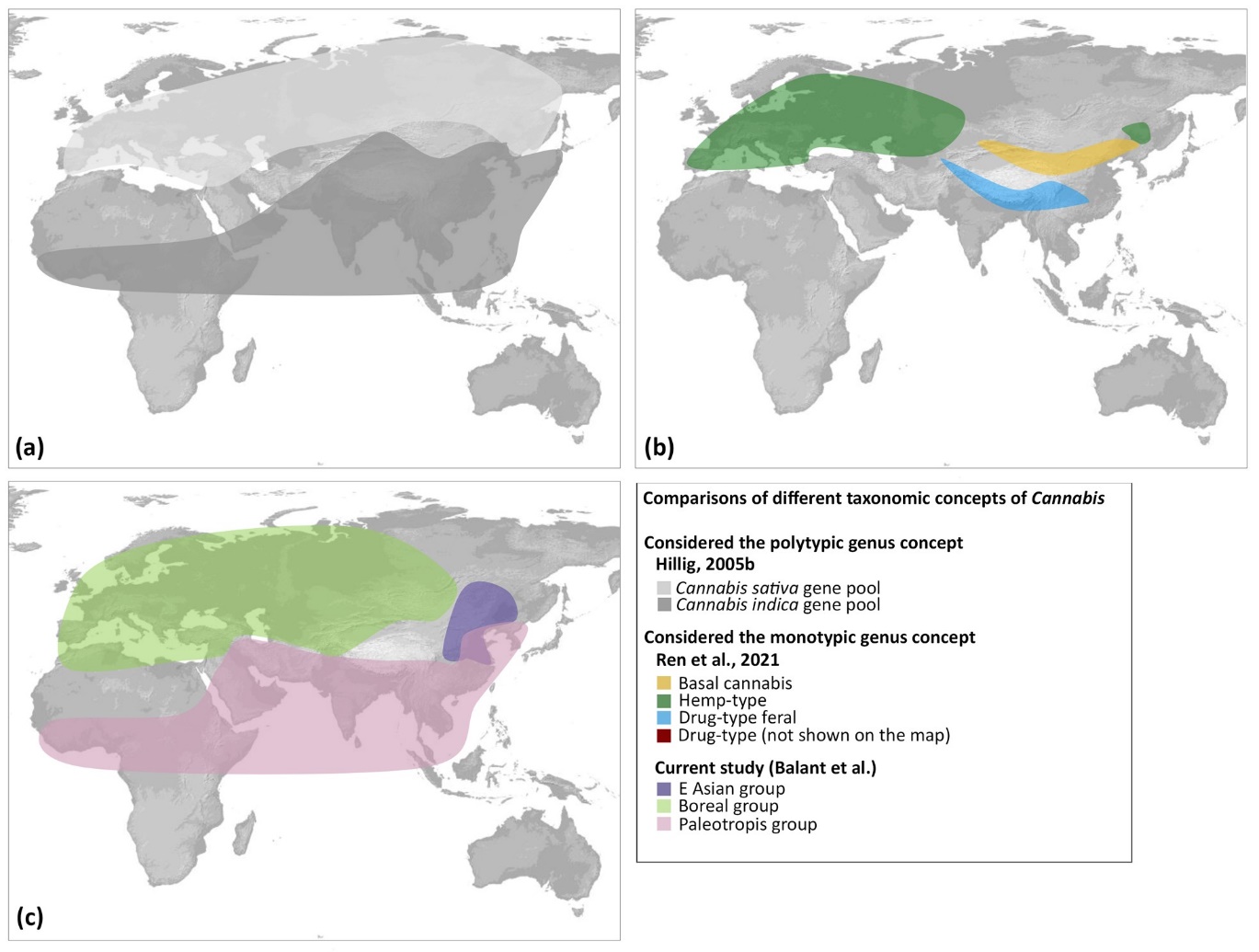

**Fig. S4** Distribution of species and genetic groups proposed by Hillig (2005b), Ren et al. (2021) and this study. (a) Hillig (2005b) considered *Cannabis* to be a polytypic genus and proposed two major gene pools (with broad ranges) and a third putative minor one (restricted to C Asia). He assigned ruderal populations from E Europe and fibre/seed landraces from Europe, Anatolia, and C Asia to the ‘*sativa* gene pool’; and feral populations from India and Nepal, fibre/seed landraces from E Asia, narrow-leafleted drug strains from S Asia, Africa, and Latin America, and wide-leafleted drug strains from Afghanistan and Pakistan to the ‘*indica* gene pool’. (b) Ren *et al.* (2021) on the other hand classified *Cannabis* as a single species, *C. sativa*, with four genetic groups: ‘Basal cannabis’, ‘Hemp-type’, ‘Drug-type’ (not shown on the map), and ‘Drug-type feral.’ The ‘Basal cannabis’ group, was found to be sister to the remaining three groups, which are named by their use type: ‘Hemp-type,’ ‘Drug-type,’ and ‘Drug-type feral’. (c) Our results support *Cannabis* as a single species (LPP = 1.0). Nonetheless, the distribution of the abovementioned ‘*sativa* gene pool’ closely matches that of our Boreal group, while the range of the ‘*indica* gene pool’ largely corresponds to our Paleotropis group.
